# Chromosome-level genome assembly of the doctor fish (*Garra rufa*)

**DOI:** 10.1101/2025.03.11.642249

**Authors:** Tetsuo Kon, Koto Kon-Nanjo, Kiki Syaputri Handayani, Liqing Zang, Fahrurrozi, Oleg Simakov, Victor David Nico Gultom, Yasuhito Shimada

## Abstract

*Garra rufa*, commonly known as doctor fish, is a small freshwater cyprinid fish recognized for its high-temperature tolerance. *G. rufa* is primarily known for its role in ichthyotherapy, in which it helps remove keratinized skin from human hands and feet. However, in recent years, its high-temperature tolerance has attracted attention owing to its potential use as a model for human disease research, including infectious diseases and cancer xenograft models. Despite this interest, the genomic basis underlying high-temperature tolerance remains largely unexplored, primarily because of the limited availability of genomic resources, which hinders its development as an experimental model. In the present study, a high-quality chromosome-level genome assembly of *G. rufa* was generated using a combination of PacBio HiFi long-read sequencing and Hi-C technology. We generated a chromosome-level genome assembly with 25 chromosomes, 1.38 Gb in total length, and scaffold N50 of 49.3 Mb. Approximately 59% of the genome comprises repetitive elements, with DNA and LTR elements being particularly abundant. In total, 27,352 protein-coding genes were annotated, of which 26,900 genes (98.3%) were functionally annotated. Benchmarking Universal Single-Copy Orthologs (BUSCO) benchmark for genome assembly and gene annotation demonstrated 94.5% and 94.7% of complete BUSCOs, respectively. We identified two heat shock transcription factor (HSF) and 239 heat shock protein (HSP)-related genes. Heat shock elements, which are HSF-binding motifs, were present within 3 kb upstream of 944 genes, with statistically significant enrichment of HSP-related genes in this set. Furthermore, molecular phylogenetic analysis and whole-genome comparisons revealed that *G. rufa* is evolutionarily closely related to species in the Labeoninae subfamily of the Cyprinidae family. This chromosome-level reference genome provides a valuable resource for future research aimed at elucidating the molecular mechanisms underlying high-temperature tolerance in *G. rufa* and establishing it as a model organism for human biomedical studies.

## Background & Summary

*Garra rufa*, commonly referred to as the doctor fish, is a small freshwater fish belonging to the cyprinid family. *G. rufa* can inhabit rivers, hot springs, and thermal pools across the Middle East, particularly in Turkey, Syria, Iraq, and Iran^1,2^. These fish are widely used in a therapeutic practice called ichthyotherapy, in which they gently nibble dead keratinized skin cells away from human hands and feet, offering a natural exfoliation treatment^3–5^. This method has gained global popularity, particularly due to its potential effectiveness in managing skin conditions like psoriasis^3–5^. Notably, ichthyotherapy sessions are conducted in hot springs or thermal pools where *G. rufa* are present. Specifically, *G. rufa* survives and exhibits feeding behavior in high-temperature environments exceeding 37°C, conditions that are challenging for the survival of other fish species.

In recent years, the growing global awareness of animal welfare and the need to reduce biomedical research costs have emphasized the importance of utilizing non-mammalian models^6,7^. Moreover, cultured cells, computational modeling, and non-mammalian models such as fish, insects, and nematodes are increasingly used^8,9^. In particular, small teleost fish are valuable for research on human diseases and drug discovery owing to their vertebrate body structures, shared organ systems with humans, and their suitability for high-throughput experiments enabled by their small size^10^. The establishment of diverse laboratory strains, development of gene manipulation tools, and progress in genome sequencing have made zebrafish (*Danio rerio*)^11–13^, medaka (*Oryzias latipes*), goldfish (*Carassius auratus*)^14–17^, and African turquoise killifish (*Nothobranchius furzeri*)^18–20^ key models for toxicology, drug discovery, and disease studies. These species share fundamental cellular mechanisms with humans, making them suitable for research in development, tissue regeneration, infectious diseases, and cancer^11–13^.

However, these conventional fish models have certain limitations, particularly regarding their optimal growth temperatures. Most small teleost fish have difficulty surviving at temperatures close to 37°C, making them less suitable for studying human diseases such as cancer and infectious diseases. For example, zebrafish, the most commonly used fish model for transplanting human-derived cancers, has a survival limit of 34°C and therefore cannot support the long-term survival of human cancer cells, as they prefer a 37°C environment^21^. In contrast, *G. rufa* can thrive in environments up to 40°C without loss of activity^22^. In addition to its high-temperature tolerance, its small body size, high fertility, ease of maintenance, and transparent eggs further enhance its potential as a model system for human disease research. Moreover, *G. rufa* belongs to the same Cyprinidae family as zebrafish, making it possible to leverage the insights gained from zebrafish studies.

Generating high-quality reference genome sequences is essential to establish *G. rufa* as an experimental model organism. Although its mitochondrial genome has been sequenced (National Center for Biotechnology Information [NCBI] accession number: NC_022941.1), the nuclear genome sequence remains unknown. Once available, the *G. rufa* genome sequence will facilitate the analysis of gene expression and regulatory mechanisms under high-temperature conditions, thereby contributing to its development as a biomedical model. Heat shock proteins (HSPs) play a critical role in protecting cells from heat-induced stress. Functioning as molecular chaperones, HSPs assist in proper protein folding and mediate the repair or degradation of damaged proteins^23–25^. Their expression is regulated by heat shock factors (HSFs), which are highly conserved transcription factors that bind to heat shock elements (HSEs) and are characterized by the consensus motif sequence nGAAnnTTCnnGAAn^23,26,27^. Indeed, a previous study demonstrated that heat stress induces the expression of HSP60, HSP70, and HSP90 in *G. rufa*, reflecting its natural thermal adaptation from river environments to hot springs^28^. However, the authors detected these *G. rufa* HSPs by western blotting using antibodies produced against human HSPs as immunogens. Therefore, high-quality chromosome-level genome assembly is essential for a deeper understanding of the HSF-HSP pathway dynamics in *G. rufa*. Beyond elucidating the molecular mechanisms underlying high-temperature tolerance, the *G. rufa* reference genome is expected to serve as a fundamental resource for genome editing, determining phylogenetic position, studying chromosome evolution, and investigating cancer xenograft-host interactions using omics approaches, such as single-cell RNA-seq. However, the genome sequences of *G. rufa* and other species belonging to the genus *Garra* have not yet been determined, hindering the progress in molecular research using *G. rufa*.

In the present study, we constructed a high-quality chromosome-level genome assembly of *G. rufa* using a combination of PacBio HiFi long-read sequencing, MGI DNBSEQ short-read sequencing, and high-throughput chromosome conformation capture (Hi-C) technology. The final genome assembly contained 25 chromosomal sequences with 1.38 Gb in total length and scaffold N50 of 49.3 Mb. In total, 27,352 protein-coding genes were annotated, of which 26,900 genes (98.3%) were functionally annotated. Approximately 58.9% of the genome comprises repetitive elements, with DNA and LTR elements being particularly prevalent. Benchmarking Universal Single-Copy Orthologs (BUSCO) analysis revealed that genome assembly and gene annotation achieved scores of 94.5% and 94.7% of complete BUSCOs, respectively. We identified two HSF genes, 239 HSP-related genes, and 1,036 putative HSEs located within the 3 kb upstream regions of 944 genes. Molecular phylogenetic analysis and whole-genome comparison revealed that *G. rufa* shares a close genetic relationship with species in the subfamily Labeoninae within the family Cyprinidae, which is consistent with the results of a previous study using mitochondrial DNA^29^. This chromosome-level reference genome provides a valuable foundation for future research on the molecular mechanisms underlying high temperature tolerance in *G. rufa* and its establishment as a model organism for human medical studies.

## Methods and Results

### Sample collection

All procedures in this study were conducted in accordance with the Japanese Animal Welfare Regulations outlined in the Act on Welfare and Management of Animals by the Ministry of Environment of Japan. Six-month-old *G. rufa* (Fig. 1a), originally collected from West Java, Indonesia, were bred at the National Research and Innovation Agency (North Lombok-West Nusa Tenggara, Indonesia). They were reared under controlled conditions at a constant temperature of 26°C, with a light cycle consisting of 12 h of light and 12 h of darkness. They were fed *ad libitum* of fish meal (Agaru, PT. Matahari Sakti, Indonesia) twice daily. Subsequently, *G. rufa* were transferred to the aquatic animal facility at Mie University, Japan, under an agreement based on the Nagoya Protocol. *G. rufa* were reared at a constant temperature of 28°C, with a light cycle of 14 h of light followed by 10 h of darkness. All experiments were conducted under anesthesia with 2-phenoxyethanol (Wako Pure Chemicals, Osaka, Japan)^30^, with every possible effort taken to reduce suffering.

**Fig. 1.**
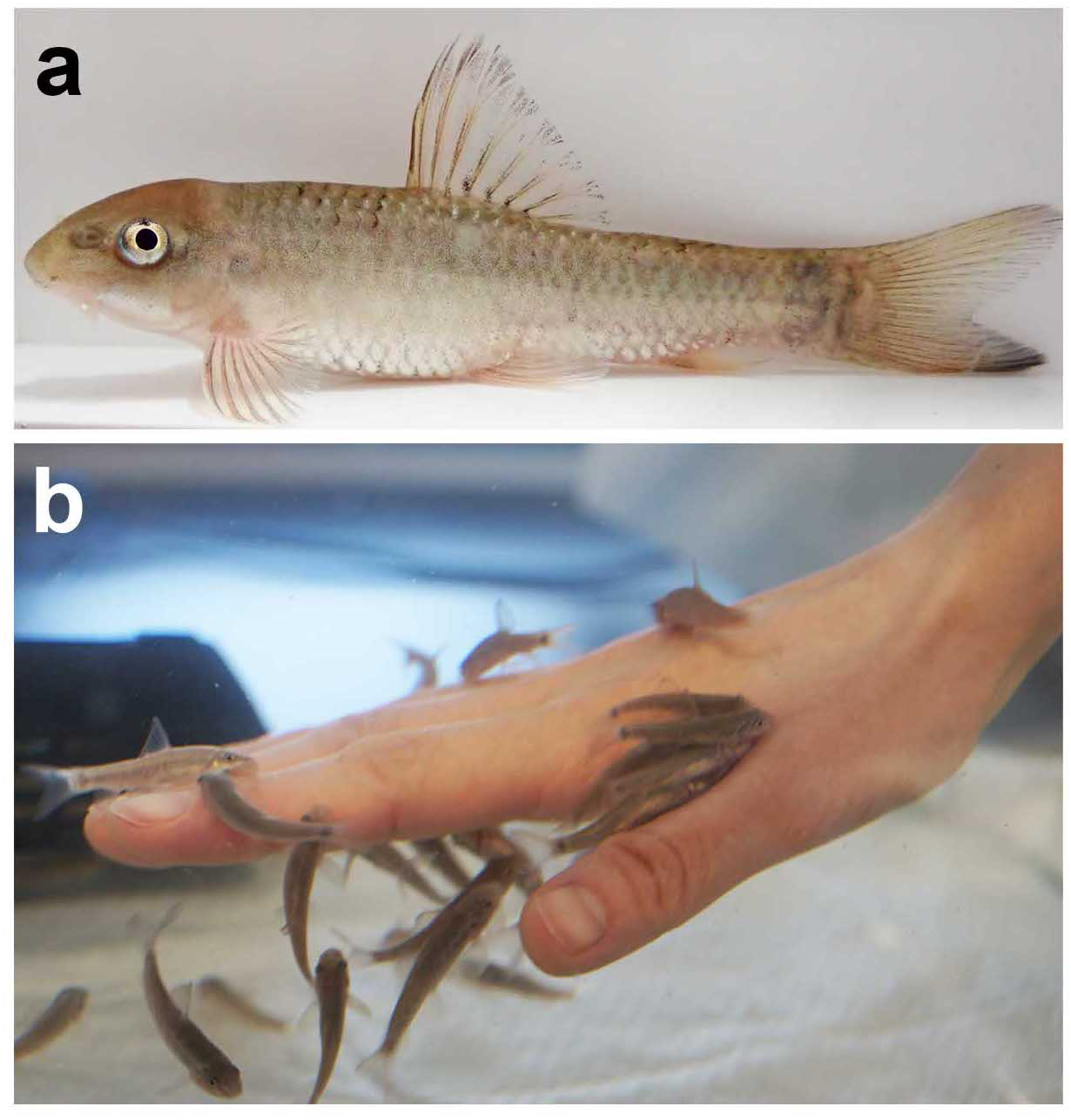
Gross appearance of *G. rufa*. (a)Lateral view of a juvenile *G. rufa*. (b)*G. rufa* individuals attached to a human hand.

To confirm the phylogenetic relationship of the sample with other species of the *Garra* genus, the cytochrome oxidase subunit-I (COI) gene sequences were amplified by PCR using the extracted genome as a template and the primers F1: 5’-TCA ACC ACC CAC AAA GAC ATT GGC AC-3’ and R1: 5’-TAG ACT TCT GGG TGG CCA AAG AAT CA-3’. The resulting PCR products were sequenced directly. The obtained sequences were aligned with publicly available 32 COI sequences from *G. rufa, G. jordanica, G. ghorensis*, and *G. nana* using Clustal Omega v1.2.4^31^ with the default parameters. The resulting multiple sequence alignments were manually curated and cropped using Jalview v2.11.4.1 (Fig. S1). Based on this sequence alignment, a neighbor-joining tree based on the Kimura 2-parameter model with gamma distribution (K2+G) was constructed with 1,000 bootstrap iterations using MEGA11^32^ (Fig. S2). The sample sequenced in the current study formed a monophyletic group with other *G. rufa* samples with 99% bootstrap support (Fig. S2). These results provide molecular phylogenetic support that our sample belongs to *G rufa*.

### Library construction and sequencing

To construct a chromosome-level genome assembly of *G. rufa*, sequencing libraries for MGI DNBSEQ short-read genome sequencing, PacBio HiFi genome sequencing, and Hi-C sequencing were constructed from a single six-month-old male (Fig. 2).

**Fig. 2.**
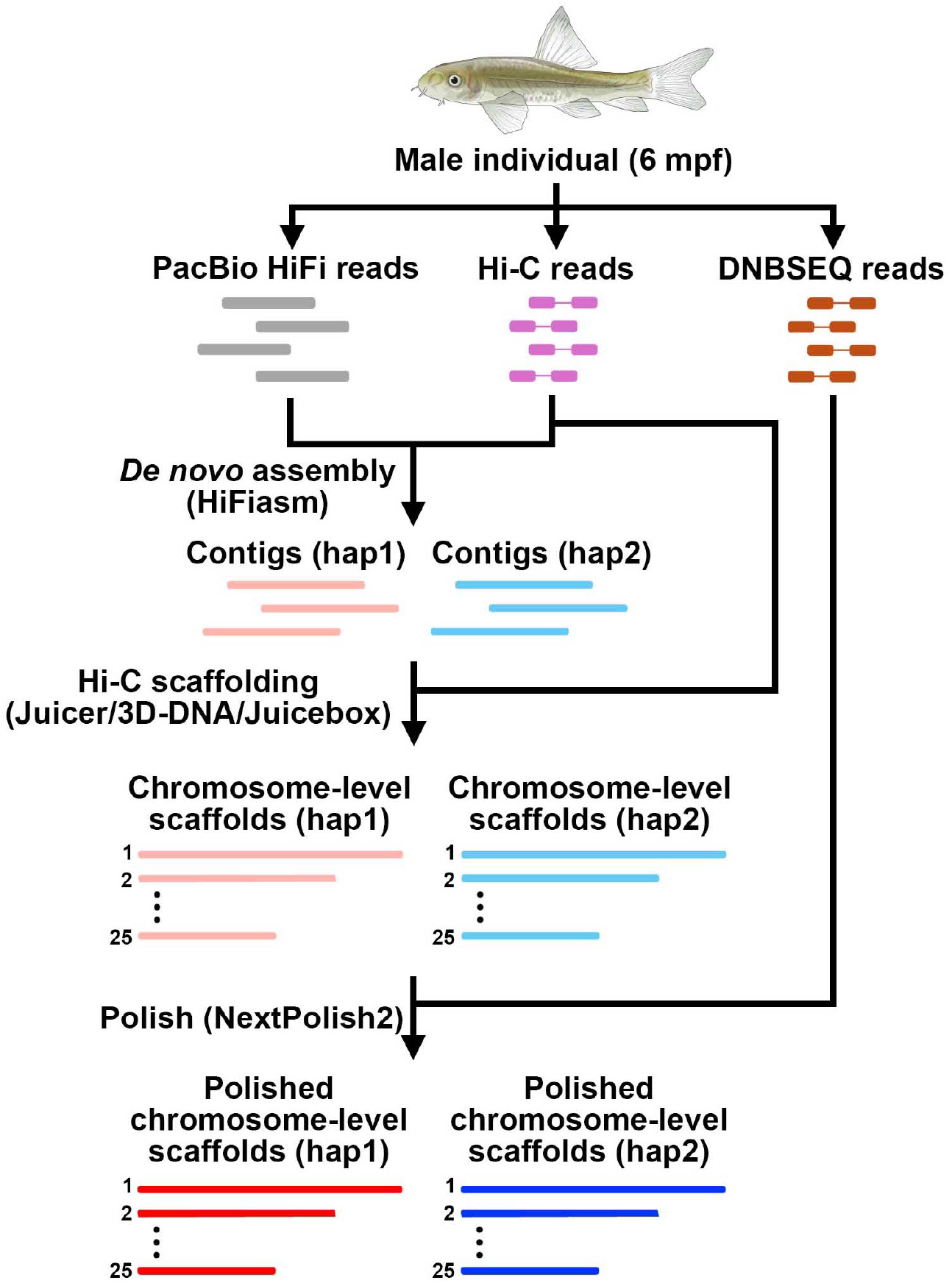
Pipeline for the chromosome-level genome assembly of *G. rufa*. The steps involved in constructing the chromosome-level genome assembly of *G. rufa* are represented in a flowchart.

For MGI DNBSEQ short-read genome sequencing, the muscle tissue was dissected from male individuals and immediately frozen in liquid nitrogen. The frozen muscle tissue was stored at −80 °C until use. Genomic DNA (gDNA) was extracted using a Lab-Aid824s DNA Extraction Kit (Zeesan Biotech, Fujian Province, China). The concentration of the extracted gDNA was measured using a Synergy LX (Agilent, CA, USA) and QuantiFluor dsDNA System (Promega, WI, USA). The size distribution of the extracted gDNA was measured using a Fragment Analyzer (Agilent, CA, USA) (Fig. S3a). MGI DNBSEQ short-read sequencing libraries were constructed using the MGIEasy Dual Barcode Circularization Kit (MGI Tech, Shenzhen, China), DNB Rapid Make Reagent Kit (MGI Tech, Shenzhen, China), and High-throughput Paired-End Sequencing Primer Kit (App-D) (MGI Tech, Shenzhen, China), following the manufacturer’s instructions. The constructed library was sequenced using the MGI DNBSEQ-T7 platform. Short-read quality was evaluated using FastQC v0.12.1^33^ (Fig. S3b). In total, 185,749,774 paired-end reads with a read length of 150 bp (55.7 Gb) were obtained (Table 1).

**Table 1.**
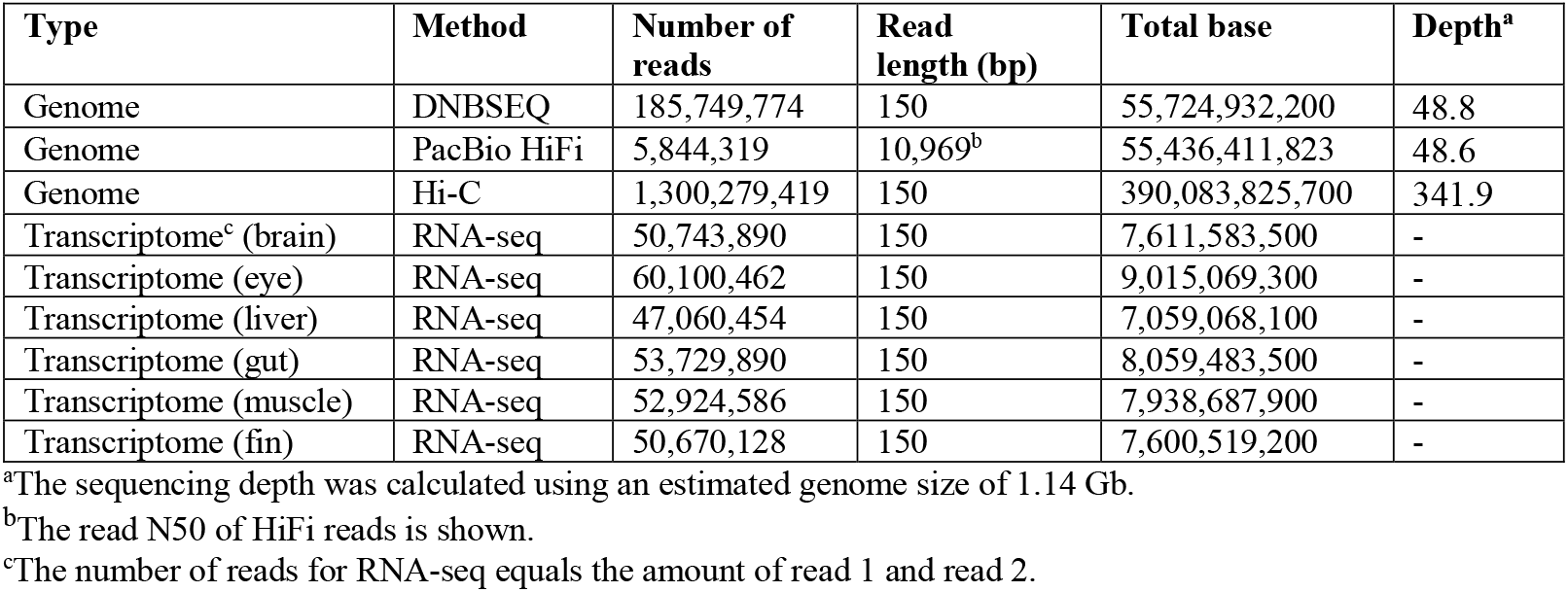
Statistics of the sequencing reads of the *G. rufa*.

| Type | Method | Number of reads | Read length (bp) | Total base | Depth<sup>a</sup> |
| --- | --- | --- | --- | --- | --- |
| Genome | DNBSEQ | 185,749,774 | 150 | 55,724,932,200 | 48.8 |
| Genome | PacBio HiFi | 5,844,319 | 10,969<sup>b</sup> | 55,436,411,823 | 48.6 |
| Genome | Hi-C | 1,300,279,419 | 150 | 390,083,825,700 | 341.9 |
| Transcriptome<sup>c</sup> (brain) | RNA-seq | 50,743,890 | 150 | 7,611,583,500 | - |
| Transcriptome (eye) | RNA-seq | 60,100,462 | 150 | 9,015,069,300 | - |
| Transcriptome (liver) | RNA-seq | 47,060,454 | 150 | 7,059,068,100 | - |
| Transcriptome (gut) | RNA-seq | 53,729,890 | 150 | 8,059,483,500 | - |
| Transcriptome (muscle) | RNA-seq | 52,924,586 | 150 | 7,938,687,900 | - |
| Transcriptome (fin) | RNA-seq | 50,670,128 | 150 | 7,600,519,200 | - |

<sup>a</sup>The sequencing depth was calculated using an estimated genome size of 1.14 Gb.
<sup>b</sup>The read N50 of HiFi reads is shown.
<sup>c</sup>The number of reads for RNA-seq equals the amount of read 1 and read 2.

For PacBio HiFi sequencing, high-molecular-weight genomic DNA (HMW gDNA) was extracted from flash-frozen muscle tissue using a genomic-tip kit (Qiagen, Venlo, Netherlands). The concentration and purity of the extracted HMW gDNA were measured using Quantus Fluorometer (Promega, WI, USA) and NanoDrop Eight Spectrophotometer (Thermo Fisher Scientific, MA, USA), respectively. The size distribution of the extracted HMW gDNA was estimated using a Fragment Analyzer (Agilent, CA, USA) to ensure that the gDNA was not fragmented (Fig. S4a and b). SMRTbell libraries were constructed following the standard protocol of PacBio (Pacific Biosciences, CA, USA), including shearing of HMW gDNA to a target size of approximately 20 kb, end repair, and hairpin adapter ligation to generate circular templates for SMRT sequencing. The SMRTbell library was sequenced using the PacBio Revio System in Circular Consensus Sequencing (CCS) mode. The quality of the PacBio HiFi reads was evaluated using NanoPlot v1.41.0 (Fig. S4c). A total of 5.84 million (55.4 Gb) Hi-Fi reads were obtained, with an N50 of 11.0 kb (Table 1).

For Hi-C sequencing, the muscle tissue was prepared from the same individual used for PacBio HiFi sequencing and immediately flash-frozen in liquid nitrogen. The DNase Hi-C (Omni-C) library was prepared using a Dovetail Omni-C Kit (Cantata Bio, CA, USA) according to the manufacturer’s instructions. Briefly, frozen tissue was ground into a fine powder using a pestle in liquid nitrogen to ensure effective crosslinking and enzyme accessibility (Fig. S5a). After chromatin crosslinking, the crosslinked chromatin undergoes nuclease digestion, fragmenting the genomic DNA while maintaining spatial proximity between interacting regions. Chromatin fragments were captured using magnetic beads that bind specifically to DNA-protein complexes. In the proximity ligation step, adjacent DNA fragments are ligated to create junctions that reflect their three-dimensional organization within the nucleus. The resulting DNA was purified and processed into sequencing libraries, which included end repairs and adapter ligation. The insert size distribution was evaluated using a Bioanalyzer (Agilent, CA, USA) (Fig. S5b). The resulting Hi-C library was sequenced using the Illumina NovaSeq X Plus sequencing system with a 150 bp read length. The quality of Hi-C reads was evaluated using FastQC v0.12.1^33^ (Fig. S5c). In total, 1,300,279,419 read pairs (390 Gb) were obtained (Table 1).

To obtain transcriptomic evidence for gene annotation, we performed RNA-Seq on six tissues (the brain, eye, liver, gut, muscle, and fin). These tissues were isolated from another six-month-old male individual by dissection using forceps (Fig. S6a–f). Total RNA was isolated using TRIzol reagent (Life Technologies, CA, USA) and a QIAGEN RNeasy Mini-prep Kit (Qiagen, Venlo, Netherlands). The RNA integrity of each sample was evaluated using agarose gel electrophoresis (Fig. S6g), and an Agilent 5400 Fragment Analyzer System (Agilent, CA, USA) (Fig. S6h). An RNA-seq library was prepared according to the manufacturer’s instructions and sequenced using the MGI DNBSEQ-T7 sequencing platform. The sequencing quality of the RNA-seq reads was evaluated using FastQC v0.12.1^33^. In total, 157,614,705 read pairs (47.2 Gb) were obtained (Table 1).

### Genome size estimation

Prior to de novo genome assembly, k-mer analysis of the MGI DNBSEQ short-reads of the *G. rufa* genome was performed to estimate genome size and heterozygosity. The frequency of the 21-mers in the MGI DNBSEQ short-reads was determined using Jellyfish v2.3.1^34^. The distribution of 21-mers showed two peaks corresponding to heterozygous and homozygous content (Fig. S7). Genome size and heterozygosity were estimated using the frequency of the 21-mers GenomeScope v2.0^35^. The size and heterozygosity of the *G. rufa* genome were estimated to be 1.14 Gb and 1.03%, respectively. Genome coverage of the repetitive elements was estimated to be 45.4%.

### *De novo* genome assembly and Hi-C scaffolding

As the initial genome assembly, we generated two haploid assemblies (haplotype 1 and haplotype 2) from the HiFi reads and Hi-C reads using HiFiasm v0.24.0-r702 ^36^ (Fig. 2). As a result, two haploid assemblies of 1.38 Gb and 1.03 Gb were obtained (Table 2). The number of contigs were 15,839 and 11,394, respectively. Contig N50 were 133.2 kb and 132.5 kb, respectively (Table 2).

**Table 2.** Genome assembly statistics.

|  | Haplotype 1 | Haplotype 2 |
| --- | --- | --- |
| Total length (bp) | 1,384,036,420 | 1,030,094,335 |
| GC content (%) | 37.0 | 36.8 |
| Contig number | 15,839 | 11,394 |
| Contig N50 (bp) | 133,182 | 132,548 |
| Contig L50 | 3,031 | 2,294 |
| Contig N90 (bp) | 40,195 | 42,023 |
| Contig L90 | 10,411 | 7,651 |
| Maximum contig length (bp) | 1,593,041 | 1,028,483 |
| Number of chromosome-level scaffolds | 25 | 25 |
| Total length of chromosome-level scaffolds | 1,301,863,782 | 993,455,113 |
| Scaffold N50 (bp) | 49,275,370 | 39,045,345 |
| Scaffold L50 | 12 | 11 |
| Scaffold N90 (bp) | 40,264,426 | 29,947,780 |
| Scaffold L90 | 24 | 22 |
| Maximum scaffold length (bp) | 108,577,553 | 87,360,613 |

Hi-C scaffolding was used to construct chromosome-level genome assemblies based on the haploid assemblies and Hi-C reads. First, the Hi-C reads were aligned to haploid assemblies and filtered using Juicer v1.6^37^. For each haploid assembly, chromosome-level scaffolds were constructed using 3D-DNA v180419^38^. The scaffolds were manually curated and assessed using Juicebox v1.11.08^39^. Finally, 25 chromosome-scale scaffolds were obtained for each haploid, where 94.1% (1.30 Gb) and 96.4% (1.03 Gb) of the contigs of the haploid assemblies were anchored to the chromosome-scale scaffolds (Fig. 3 and S8). This result aligns with that of a previous karyotyping study, which reported that *G. rufa* has 2n = 50 chromosomes^1^. The scaffold N50 values were 49.3 Mb and 39.0 Mb for each haploid genome, respectively (Table 2). To polish these two haploid chromosome-level genome assemblies, the MGI DNBSEQ short-reads were trimmed using Trimmomatic v0.39^40^ using the parameters “PE -phred33 ILLUMINACLIP:adapters.fa:2:30:10 TRAILING:30 LEADING:30 SLIDINGWINDOW:4:25 MINLEN:50.” Using the trimmed DNBSEQ short-reads, genome assemblies were polished using NextPolish2 v0.2.1-4fec66b^41^.

**Fig. 3.**
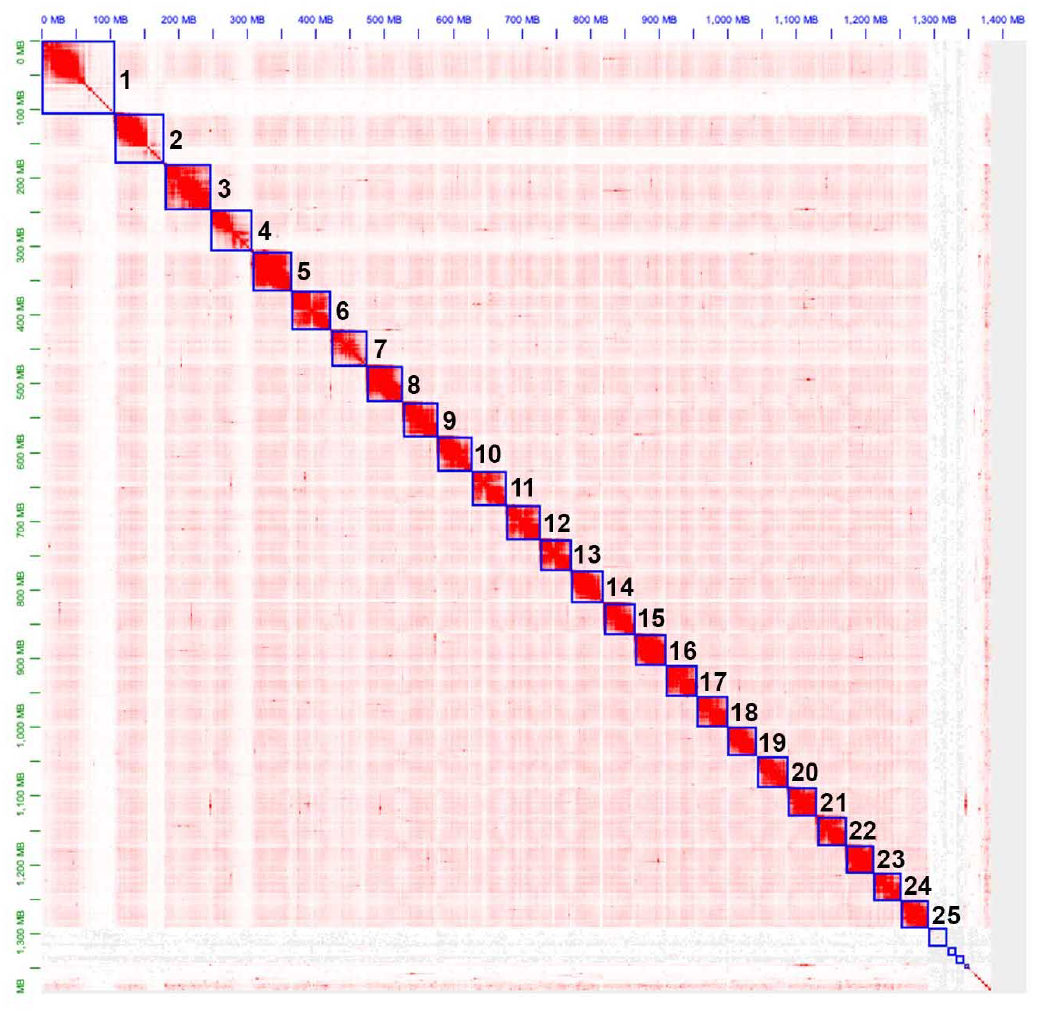
Chromosome-level genome assembly of *G. rufa*. Hi-C contact map for the haplotype 1 is shown. Each blue square represents the boundary between the scaffolds. Chromatin contact signals are shown as red spots, with their intensity reflecting the degree of chromatin contact strength. There are 25 chromosome-scale scaffolds.

The completeness of the genome assemblies, we obtained the Actinopterygii odb10 database (actinopterygii_odb10) using BUSCO software v5.8.2^42^ with the option “--download actinopterygii_odb10.” Then, BUSCO benchmark analysis was performed on both of the haplotype genome assemblies with options “-m genome.” In the haplotype 1 genome, out of a total of 3,640 BUSCOs, we identified 3,440 (94.5%) complete BUSCOs, including 3,142 (86.3%) complete and single-copy BUSCOs and 298 (8.2%) complete and duplicated BUSCOs. The numbers of fragmented and missing BUSCOs were 77 (2.1%) and 123 (3.4%), respectively (Table 3). Similarly, in the haplotype 2 genome, there were 3,013 (82.8%) complete BUSCOs, comprising 2,888 (79.3%) complete and single-copy BUSCOs, and 125 (3.4%) complete and duplicated BUSCOs. The numbers of fragmented and missing BUSCOs were 116 (3.2%) and 511 (14.0%), respectively (Table 3).

**Table 3.** Completeness of the *G. rufa* genome assembly and its gene annotation.

|  | Genome (Haplotype 1) |  | Genome (Haplotype 2) |  | Gene annotation (Haplotype 1) |  |
| --- | --- | --- | --- | --- | --- | --- |
|  | Number | Percentage (%) | Number | Percentage (%) | Number | Percentage (%) |
| Complete BUSCOs (C) | 3,440 | 94.5 | 3,013 | 82.8 | 3,446 | 94.7 |
| Complete and single-copy BUSCOs (S) | 3,142 | 86.3 | 2,888 | 79.3 | 3,209 | 88.2 |
| Complete and duplicated BUSCOs (D) | 298 | 8.2 | 125 | 3.4 | 237 | 6.5 |
| Fragmented BUSCOs (F) | 77 | 2.1 | 116 | 3.2 | 19 | 0.5 |
| Missing BUSCOs (M) | 123 | 3.4 | 511 | 14.0 | 175 | 4.8 |
| Total BUSCOs searched | 3,640 | 100 | 3,640 | 100 | 3,640 | 100 |

Thus, we successfully obtained a haplotype-resolved chromosome-scale genome assembly of *G. rufa*. For downstream analyses, we used the genome assembly of haplotype 1, which exhibited higher scaffold N50 and BUSCO scores than those of haplotype 2 (Table 3).

### Mitochondrial genomic DNA

The assembled contigs were analyzed using BLASTN v2.16.0^43^ to search for sequences homologous to the mitochondrial genomic sequence of *G. rufa* (NC_022941.1, 16,763 bp) available in the NCBI nucleotide database. A single hit was identified, which was 16,760 bp in length and formed an alignment across the entire query length (Fig. S9a). The sequence identity between the two sequences was 99.24%. This contig of the mitochondrial genome was extracted and annotated using MitoAnnotator Web Server^44^. Consequently, 13 protein-coding genes, two ribosomal RNA genes, and 22 transfer RNA genes were identified (Fig. S9b).

### Analysis of repetitive elements in the *G. rufa* genome

To identify repetitive elements in the *G. rufa* genome, a custom repeat library was generated using RepeatModeler v2.0.5^45^ with the default parameters. Unclassified repeat sequences were further annotated by searching against the Dfam 3.8 database^46^ using nhmmscan v3.4^47^ to improve the repetitive element annotation^48^. RepeatMasker v4.1.7-p1^49^ was used to detect repetitive elements in the genome assembly to produce a soft-masked genome sequence. A total of 820 Mb (58.9%) of the genome sequence was occupied by repetitive elements. Among the repetitive elements, SINEs, LINEs, LTRs, and DNA transposons constituted 2.8%, 4.8%, 9.1%, and 34.4% of the total repeat sequences, respectively (Fig. 4a; Table 4). Furthermore, the distribution of repetitive elements was analyzed separately for chromosome-level scaffolds and unplaced sequences. The results showed that 57.2% of the chromosome-level scaffolds were occupied by repetitive elements, whereas 83.5% of the unplaced sequences consisted of repetitive elements. Notably, the proportion of LTR elements was higher in unplaced sequences than in chromosome-level scaffolds, suggesting that their repetitive nature may explain the difficulty of anchoring these sequences to chromosomes (Fig. 4b,c; Table 4).

**Table 4.** Repetitive elements in the G. *rufa* genome assembly.

| Repetitive element type | All |  |  | Chromosomes |  |  | Unplaced sequences |  |  |
| --- | --- | --- | --- | --- | --- | --- | --- | --- | --- |
|  | Number of elements | Length (bp) | Genome coverage (%) | Number of elements | Length (bp) | Genome coverage (%) | Number of elements | Length (bp) | Genome coverage (%) |
| SINE | 154,204 | 39,087,872 | 2.8 | 136,663 | 34,957,409 | 2.7 | 17,542 | 4,130,776 | 4.61 |
| LINE | 259,976 | 66,328,278 | 4.8 | 244,671 | 60,013,228 | 4.6 | 15,313 | 6,316,041 | 7.1 |
| LTR | 196,338 | 126,703,625 | 9.1 | 183,755 | 100,701,704 | 7.7 | 12,587 | 26,002,426 | 29.0 |
| DNA | 2,648,231 | 478,038,547 | 34.4 | 2,565,430 | 455,176,539 | 35.0 | 82,869 | 22,878,006 | 25.5 |
| RC | 153,825 | 71,798,606 | 5.2 | 140,182 | 57,717,337 | 4.4 | 13,661 | 14,090,344 | 15.7 |

**Fig. 4.**
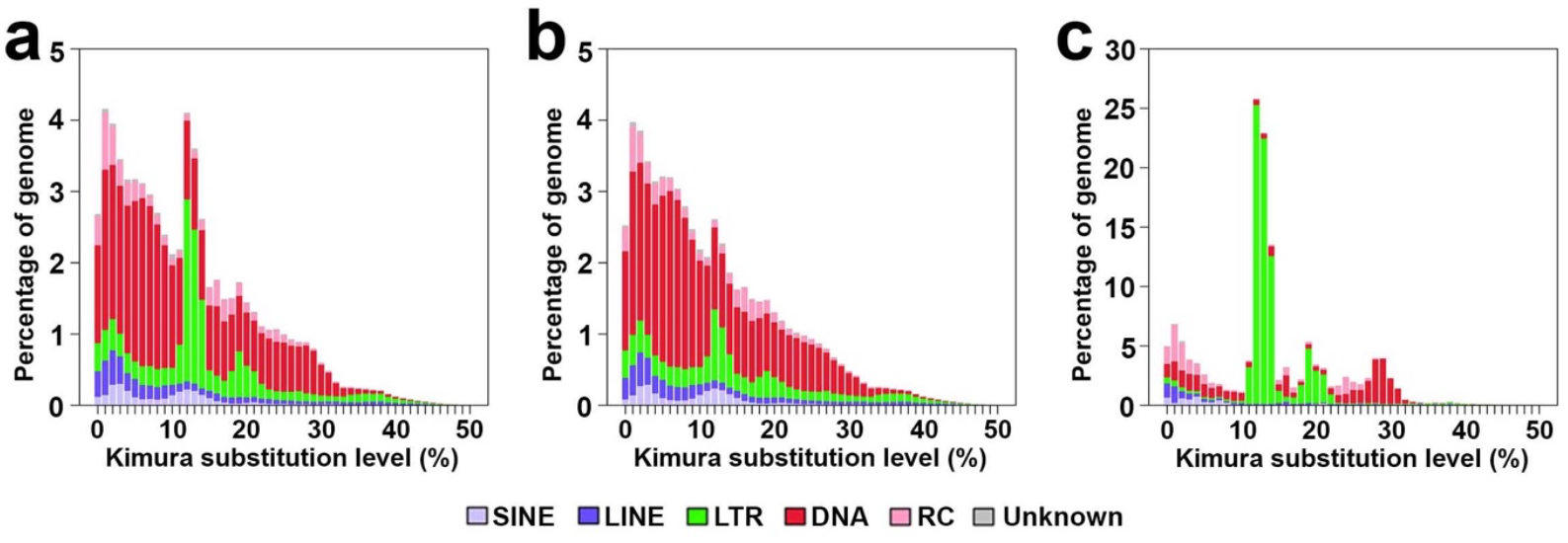
Landscape of repetitive elements in the *G. rufa* genome assembly. Distribution of repetitive elements in the whole-genome assembly (a), chromosome-scale scaffolds (b), and unplaced sequences (c). The x-axis represents the Kimura substitution level (%), which estimates the evolutionary divergence of the repetitive elements from their consensus sequences. The y-axis shows the percentage of the genome, indicating the relative abundance of each type of repetitive element at different divergence levels.

### Gene prediction and functional annotation

To predict genes in the *G. rufa* genome assembly, we used a integrative approach combining transcriptome-based, de novo, and homology-based methods using BRAKER3^50^. To obtain transcriptome evidence, the RNA-seq reads from the six tissues were aligned to the soft-masked genome assembly using HISAT2 v2.2.1^51^ with the default parameters. For evidence of protein-homology, we used 1,444,243 protein sequences 24 Cypriniformes species (Table S1). The singularity image of BRAKER3^52^ was used for the genome annotation. BRAKER3 was run on the soft-masked genome with a parameter “--softmasking --gff3”. Consequently, we predicted 27,352 protein-coding genes in the genome. These genes have an average coding sequence length of 549 bp, an average gene length of approximately 15.9 kb, and contain an average of 8.6 exons. These genes were widely distributed across all the chromosomes (Fig. 5a).

**Fig. 5.**
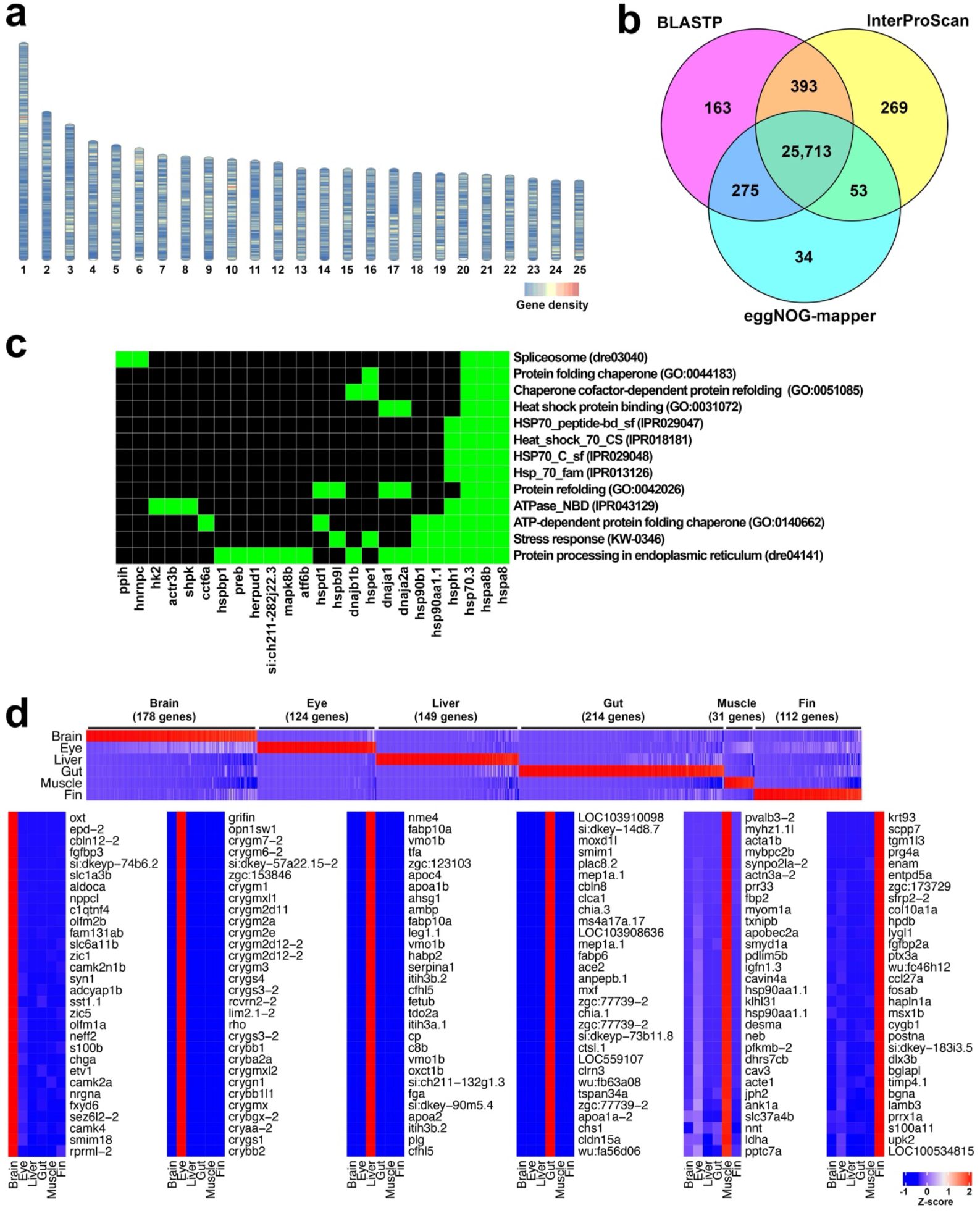
Gene functional annotation and gene expression profiles. (a) Gene density across chromosomes. (b) Venn diagram showing the results of functional gene annotation. (c) Top annotation cluster of DAVID analysis on the 944 genes with heat shock elements within 3 kb upstream regions. (d) Gene expression profiles. The upper panel displays the expression profiles of 808 genes showing tissue-specific expression. The bottom panels show the top 30 genes for each of the six tissues, ranked by fold change calculated as the expression level in a given tissue divided by the average expression level across other tissues. The transcripts per million value of each gene is normalized using Z-transformation.

To assess the completeness of the gene prediction, protein sequences were generated from the genome sequence and genome annotation using GffRead v0.12.7^53^, with the longest protein isoform extracted from each gene. These protein sequence sets were subjected to BUSCO benchmark analysis ^42^. Of the total 3,640 BUSCOs, we identified 3,446 complete BUSCOs (94.7%), including 3,209 (88.2%) single-copy and 237 (6.5%) duplicated BUSCOs (Table 3). The numbers of fragmented and missing BUSCOs were 19 (0.5%) and 175 (4.8%), respectively (Table 3).

The predicted gene sets were further analyzed using functional annotations. We used BLASTP v2.16.0^43^ with the parameter “-evalue 1e-20” to search the 27,352 protein sequences from the predicted genes against the 1,444,243 protein sequences from the 24 Cypriniformes species used for genome annotation (Table 5,S1). In total, 26,544 (97.0%) predicted proteins had hits in this search (Table 5,S2). A total of 26,428 (96.6%) predicted proteins were functionally annotated using InterProScan v5.72-103.0^54^ (Table 5). Also, gene functional annotation was conducted using eggNOG-mapper v2.1.12^55^ and 26,075 (95.3%) of the predicted proteins had hits (Table 5,S3). Overall, a total of 26,900 (98.3%) genes were assigned to one or more functional annotation (Fig. 5b; Table 5,S4). Furthermore, protein sequences of 25,167 genes (92.0%) out of the 27,352 predicted genes corresponded to in zebrafish genes based on BLASTP v2.16.0^43^ search with the parameter “-evalue 1e-20” (Table S5).

**Table 5.** Functional annotation of the protein-coding genes in the *G. rufa* genome.

| Category | Number | Percentage (%) |
| --- | --- | --- |
| Total genes | 27,352 | 100 |
| BLASTP against proteins of Cypriniformes | 26,544 | 97.0 |
| InterProScan | 26,428 | 96.6 |
| eggNOG-mapper | 26,075 | 95.3 |
| Total annotated genes | 26,900 | 98.3 |

### HSP-related genes in the *G. rufa* genome

A previous study showed that heat stress trigged the expression of HSP60, HSP70, and HSP90 in *G. rufa*, suggesting a potential contribution of HSPs to the high-temperature tolerance^28^. We confirmed that *G. rufa* possesses two paralogs of HIF-1α (GR02496, GR15074), reported in zebrafish^56^. Furthermore, when we performed a BLASTP search using proteins encoded by *G. rufa* genes against 90 human HSP-related proteins^57^, including HSP60, HSP70, and HSP90, we found that 239 *G. rufa* genes matched 75 human HSP-related genes (Table S6). A similar analysis in zebrafish revealed that 231 zebrafish genes matched 76 HSP-related genes (Table S7). Furthermore, to identify a set of genes that could potentially be regulated by HIF, we searched for genes containing the HSE motif sequence (nGAAnnTTCnnGAAn)^58^ within 3 kb upstream of the gene using the fuzznuc tool^59^. As a result, we identified a total 1,036 HSEs within the 3kb-upstream regions of 944 genes, among which 16 were HSP-related genes (Table S8), representing a statistically significant enrichment (P < 0.05, Fisher’s exact test). Furthermore, functional annotation clustering of the 944 genes was performed using the DAVID web server^60^. Three significantly enriched annotation clusters (enrichment score > 1.3) were identified. The top annotation cluster (enrichment score: 2.14) included annotation terms such as “heat shock protein binding (GO Biological processes, GO:0031072)”, “protein refolding (GO Biological processes, GO:0042026)”, and “Protein processing in endoplasmic reticulum (KEGG passway, dre04141)” (Fig. 5c, Table S9). These observations highlight that the genome assembly of *G. rufa* and its gene annotation serve as essential resources for future comprehensive studies on HIF-mediated regulation of HSP-related genes in this species.

### Gene expression profiling of the six tissues

We verified whether gene expression could be appropriately measured based on the gene annotations. RNA-seq reads from the six tissues for genome annotation were mapped to the genome assembly using HISAT2 v2.2.1^51^ with the default parameters. Then, featureCounts v2.0.8^61^ was used to count the number of reads mapped to each predicted gene, and transcripts per million (TPM) values were calculated (Table S10). To identify genes with tissue-specific expression, we extracted genes with TPM values greater than 200 from a given tissue and an average expression level of 100 or less across five other tissues. We identified 808 genes with tissue-specific expressions (Fig. 5c; Table S11). Rhodopsin and crystallin genes were expressed in eye samples, consistent with their known roles in rod photoreceptor cells and the lens, respectively. Similarly, actin alpha 1, skeletal muscle (acta1) was highly expressed in muscle samples (Fig. 5d). These observations demonstrate that the gene expression patterns align with the expected biological roles, confirming that our approach effectively captures tissue-specific expression and supports the reliability of gene annotations.

### Molecular phylogenetic analysis

In previous studies, morphological and molecular phylogenetic analyses using several gene sequences suggested that *G. rufa* belongs to the Labeoninae subfamily^62,63^. However, its phylogenetic position has not been examined based on a large gene set. Therefore, we analyzed the phylogenetic relationships between *G. rufa* and 12 representative Cypriniformes species, whose chromosome-scale genome assemblies have been determined (Table S12). To extract single-copy orthologs for molecular phylogenetic analysis in an unbiased manner, we selected species with 24 or 25 chromosomes that were presumed not to have undergone lineage-specific whole-genome duplication^15,16^. The genome sequences and annotations of these species were obtained from the NCBI genome database. Protein sequences were retrieved from these genome sequences and annotations using GffRead v0.12.7^53^. The longest protein isoform of each gene was used for further analyses. To identify gene families, OrthoFinder v3.0.1b1^64^ was performed on these protein sets. In total, 20,340 orthogroups were identified. Among them, 10,793 single-copy orthogroups were identified (Table S13). Protein sequences within each of the 10,793 single-copy orthogroup were aligned using MAFFT v7.526^65^. All protein sequence alignments were concatenated into a single protein sequence alignment with 6,897,643 positions. The alignment were then refined using GBLOCKS v0.91b^66^ with parameters “-t=p -b1=7 -b2=7 - b3=8 -b4=10 -b5=n” to remove poorly aligned and ambiguously aligned regions, resulting in a final alignment of 5,427,445 positions (78% of the original dataset). For model selection, ProtTest3 v 3.4.2^67^ was used to determine the best-fit amino acid substitution model based on the Akaike information criterion. The JTT+I+G model was identified as the best-fit model. This model was subsequently applied for maximum likelihood (ML) phylogenetic inference using RAxML v8.2.12^68^. The ML tree was constructed with 1,000 bootstrap replicates to assess branch support, and the final tree was visualized using MEGA11^69^. In the resulting ML tree, *G. rufa* formed a monophyletic group with *Labeo rohita*, a Labeoninae species, with 100% bootstrap support, confirming that *G. rufa* belongs to the Labeoninae subfamily (Fig. 6a). A NJ tree was also constructed with 1,000 bootstrap replicates. In this tree, *G. rufa* also formed a monophyletic group with *L. rohita* with 100% bootstrap support (Fig. S10).

**Fig. 6.**
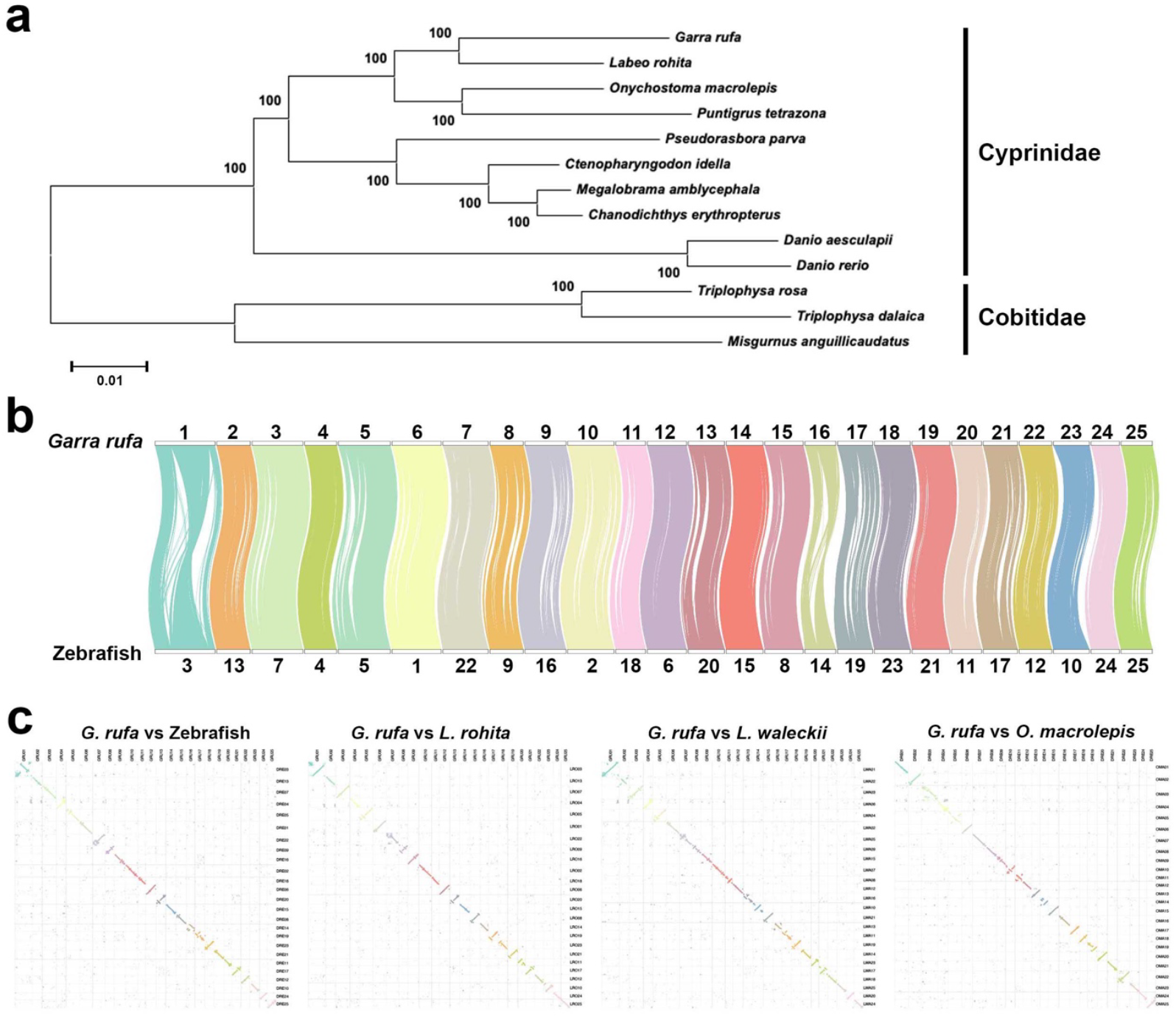
Molecular phylogenetic analysis of *G. rufa* and other representative Cyprinidae species. (a) Maximum likelihood tree *G. rufa* and Cypriniformes species with 1,000 bootstrap iterations. The scale bar represents a phylogenetic distance of 0.01 amino acid substitutions per site. (b) Chord plot showing the chromosomal synteny between *G. rufa* and zebrafish (*D. rerio*). The curved lines connecting the chromosomes indicate orthologs that form linkage groups. (c) Oxford dot plot showing chromosomal synteny between two species. Each dot represents a gene, and the coordinates are based on the gene index on their respective genomes.

### Chromosomal synteny analysis

To investigate chromosome-scale conservation, we analyzed chromosomal homology. Using the R package MacrosyntR^70^, we extracted genes forming linkage groups^71–73^ between *G. rufa* and zebrafish. Both the chord plot (Fig. 6b) and theoxford dot plot (Fig. 6c) revealed that the chromosomal collinearity between *G. rufa* and zebrafish was highly conserved. Furthermore, we detected a high degree of chromosomal collinearity between G. rufa and each of *Labeo rohita, Leuciscus waleckii*, and *Onychostoma macrolepis* (Fig. 6c).

### Data Records

All raw sequencing data produced in this study, including PacBio HiFi sequencing, HiC, and RNA-seq, have been deposited in the NCBI SRA database under the BioProject accession number PRJNA1204062. The MGI DNBSEQ reads were deposited under the accession number SRR32173939. The PacBio HiFi reads of the genome were deposited under the accession number SRR31966927. The Hi-C reads were archived under the accession number SRR31846638. The RNA-seq reads of the six tissues were archived under the accession number SRR32181860–SRR32181865. The genome assembly was deposited in the NCBI Genome database under the accession number JBLIWC000000000.

### Technical Validation

The quality of the gDNA and RNA was evaluated in terms of purity, concentration, and size distribution to ensure high quality before sequencing. The quality of the PacBio HiFi reads was assessed using NanoPlot v 1.40.2^74^, whereas the quality of the DNBSEQ, Hi-C, and RNA-seq reads was evaluated using FastQC v0.12.1^33^ before downstream analysis. To evaluate the genome assembly quality, the PacBio HiFi reads of the *G. rufa* genome were mapped back to the genome assembly using Minimap2 v2.26-r1175^75^ with option “-ax map-hifi”. A total of 99.8% of the PacBio HiFi reads were mapped. Similarly, mapping the DNBSEQ short reads to the genome assembly using BWA-MEM v0.7.13-r1126^76^ resulted in 98.3% being mapped. The RNA-seq reads from the six samples used for genome annotation were aligned to the genome using HISAT2 v2.2.1^51^, resulting in an average mapping rate of 92.9%. The completeness of the genome assembly and gene annotation was evaluated using BUSCO v5.8.2 and the Actinopterygii odb10 database (actinopterygii_odb10). The BUSCO scores for the genome assembly and gene annotation were 94.5% and 94.7%, respectively.

## Code availability

No custom codes were developed for this study. All software was used according to the manuals and protocols provided by the software developers. If no detailed parameters were mentioned for the software, it was run using its default parameter settings, as described in the software documentation.

## Acknowledgments

The authors thank Ms. Kyoko Shimada at Mie University and Dr. Herbert Gasser at the University of Vienna for their support in organizing this study. This work was supported by the Okasan-Katoh Foundation, Takahashi Industrial and Economic Research Foundation and JSPS KAKENHI (grant number 24K21914) to YS, Takeda Science Foundation to TK, Mochida Memorial Foundation for Medical and Pharmaceutical Research to TK, International Medical Research Foundation to TK, Yamada Science Foundation to TK, Austrian Science Fund FWF (I4353), and ERC-H2020-EURIP grant (grant number 945026) to OS. Computations were carried out on the Life Science Compute Cluster (LiSC) of the University of Vienna.

## Contributions

T.K., O.S., and Y.S. conceived and designed the study. K.H., F., V.G., and Y.S. participated in sample preparation. T.K., K.KN., K.H., V.G., and Y.S. performed the HiC experiments and genome sequencing. L.Z. and Y.S. performed RNA-seq experiments. T.K., K.KN., and O.S. performed bioinformatic analyses. K.H., V.G., and Y.S. performed mitochondrial genome sequencing. T.K., K.KN., and Y.S. wrote and revised the manuscript with comments from all the authors. All the authors have read and approved the final version of the manuscript.

## Ethics declarations

### Conflicts of interest

The authors declare no competing interests.

## Supplementary figures

Fig. S1. Multiple sequence alignment of the cytochrome oxidase subunit-I (COI) genes from *Garra* species

Fig. S2. Neighbor-joining tree of *G. rufa, G. jordanica, G. ghorensis*, and *G. nana* constructed using the COI sequence alignment

Fig. S3. Sequencing quality of the MGI DNBSEQ short-read sequencing of the genome Fig. S4. Size distribution of the extracted high-molecular-weight genomic DNA

Fig. S5. Hi-C sequencing of the *G. rufa* genome

Fig. S6. Total RNA extraction from *G. rufa* tissues for RNA-seq analysis

Fig. S7. The k-mer distribution of the Illumina genome sequencing reads using GenomeScope based on a k value of 21

Fig. S8. Hi-C contact map of the haplotype 2 Fig. S9. Mitochondrial genome of *G. rufa*

Fig. S10. Neighbor-joining tree showing the relationship between *G. rufa* and other cypriniforms

## Supplementary tables

Table S1. Species used to generate the genome annotation of *G. rufa*

Table S2. Results of the BLASTP search of predicted proteins against the proteins of 24 Cypriniformes species

Table S3. Functional annotation using EggNOG-mapper Table S4. List of functionally annotated genes

Table S5. Results of the BLASTP search of predicted proteins against the zebrafish proteins Table S6. Heat shock protein (HSP)-related genes of *G. rufa*

Table S7. HSP-related genes of zebrafish

Table S8. 1,036 Heat shock elements (HSEs) located within 3kb-upstream regions of 944 genes

Table S9. Functional annotation clustering of the 944 genes that have HSEs within 3 kb upstream

Table S10. Gene expression profile

Table S11. List of genes showing tissue-specific expression Table S12. Species used for the molecular phylogenetic analysis

Table S13. Single-copy orthogroups among *G. rufa* and 12 representative Cypriniformes species

## Notes

### Competing Interest Statement

The authors have declared no competing interest.

