## Supplementary figures for "Chromosome-level genome assembly of the doctor fish (*Garra rufa*)"

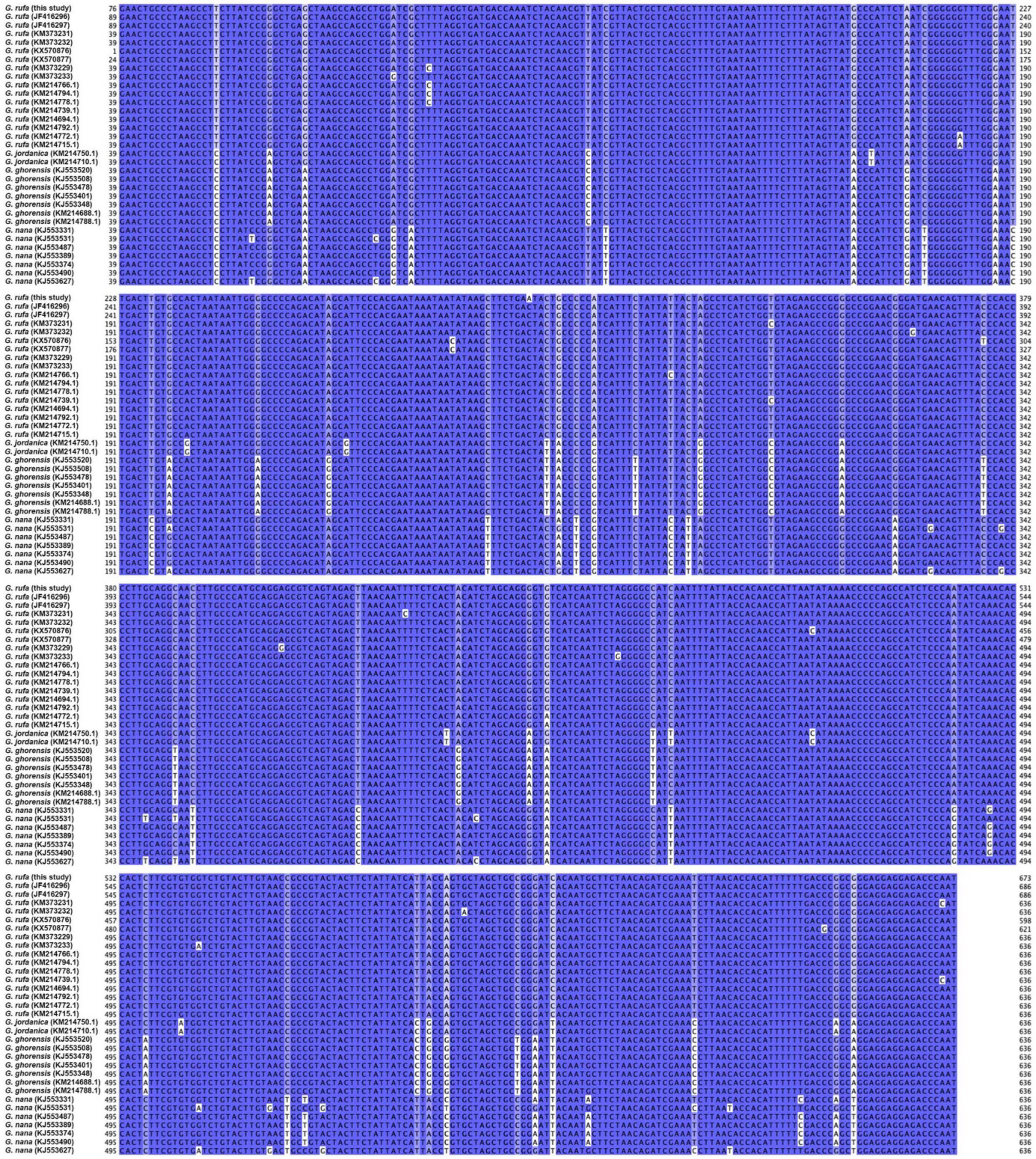

**Fig. S1. Multiple sequence alignment of the cytochrome oxidase subunit-I (COI) genes from *Garra* species**

Each sequence name on the left side of the alignment indicates the species name and GenBank accession number. Each base position is colored according to the degree of sequence identity.

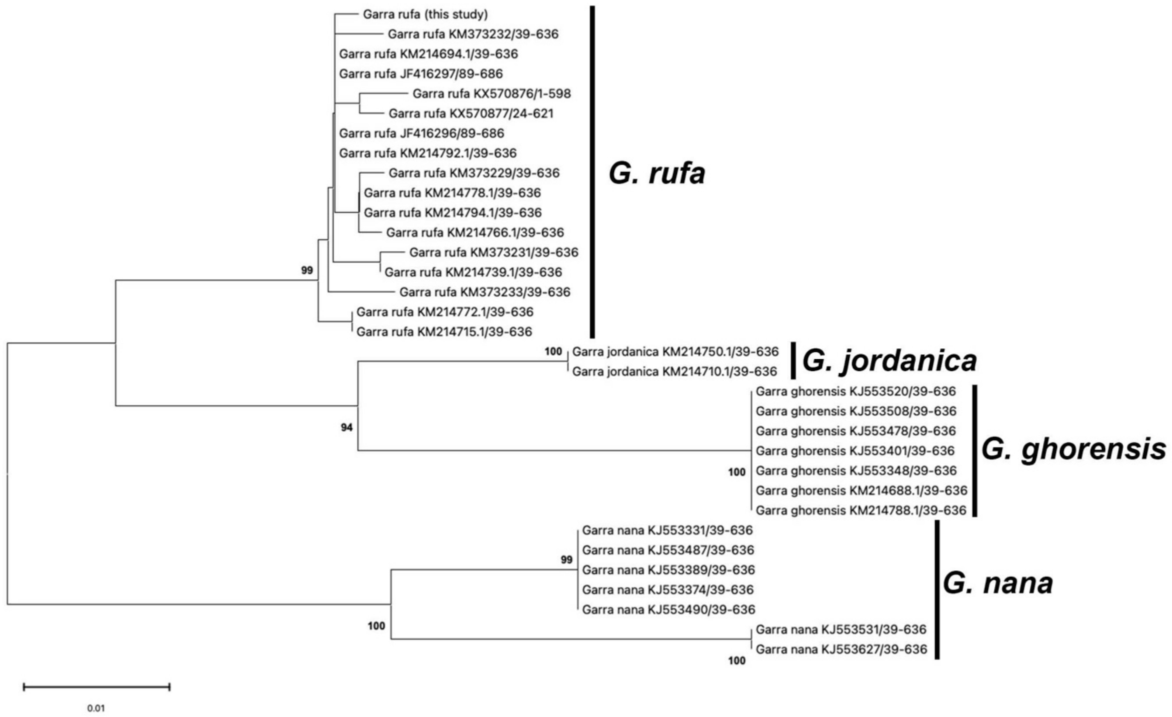


**Fig. S2. Neighbor-joining tree of *G. rufa*, *G. jordanica*, *G. ghorensis*, and *G. nana* constructed using the COI sequence alignment**

A neighbor-joining tree of *Garra* species was constructed based on the Kimura 2-parameter model with gamma distribution (K2+G) and 1,000 bootstrap replicates. Bootstrap values exceeding 90% are indicated. The scale bar represents the phylogenetic distance of 0.01 nucleotide substitutions per site. The samples sequenced in this study formed a monophyletic group with the other *G. rufa* samples, supported by a bootstrap value of 99%.

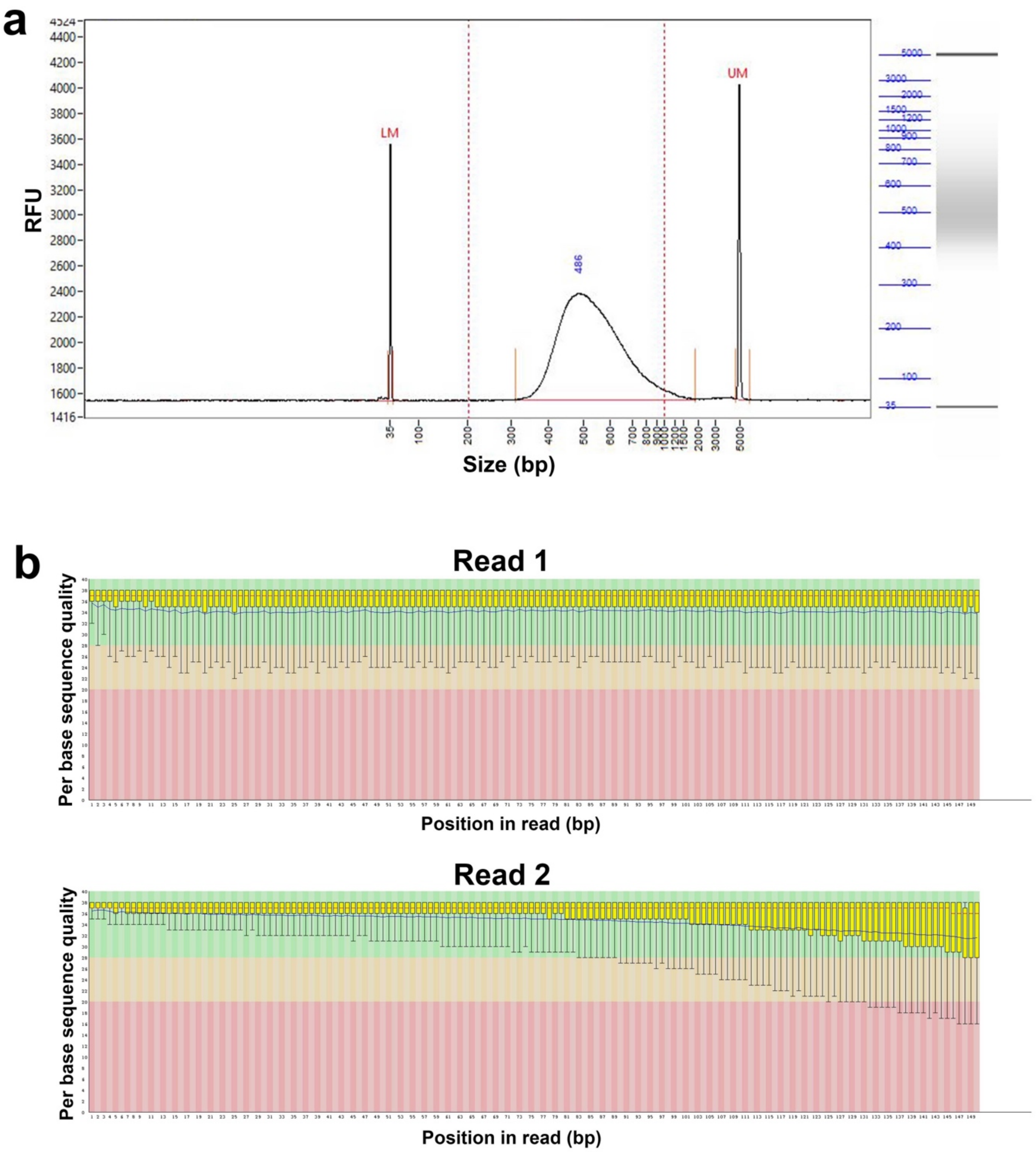


**Fig. S3. Sequencing quality of the MGI DNBSEQ short-read sequencing of the genome**
(a) Insert size distribution of the MGI DNBSEQ short-read genome sequencing library using the Fragment Analyzer (Agilent, CA, USA). Size distribution is shown in the electropherogram (left panel) and gel image (right panel). Abbreviations: LM, lower marker; UM, upper marker; RFU, relative fluorescence unit. (B) Per-base sequencing quality of MGI DNBSEQ short-read genome sequencing. The figure was generated using the FastQC program.

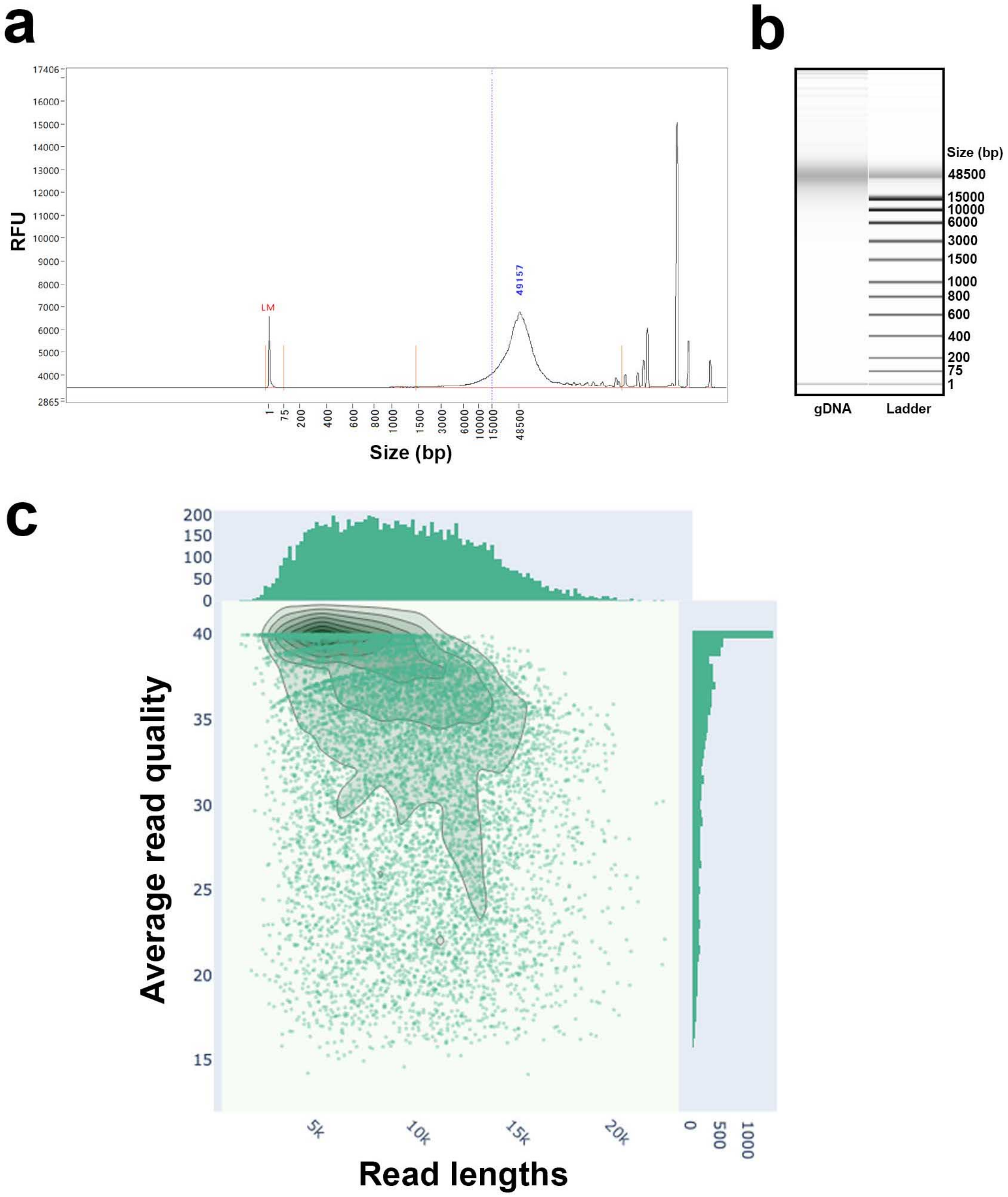


**Fig. S4. Size distribution of the extracted high-molecular-weight genomic DNA**

(a, b) Size distribution of the extracted genome DNA was analyzed using the Fragment Analyzer (Agilent, CA, USA). The result is presented in the electropherogram (a) and gel image (b). Abbreviations: LM, lower marker; RFU, relative fluorescence unit. (c) Read length and average read quality of the PacBio HiFi long-reads. The solid lines represent kernel density estimation.

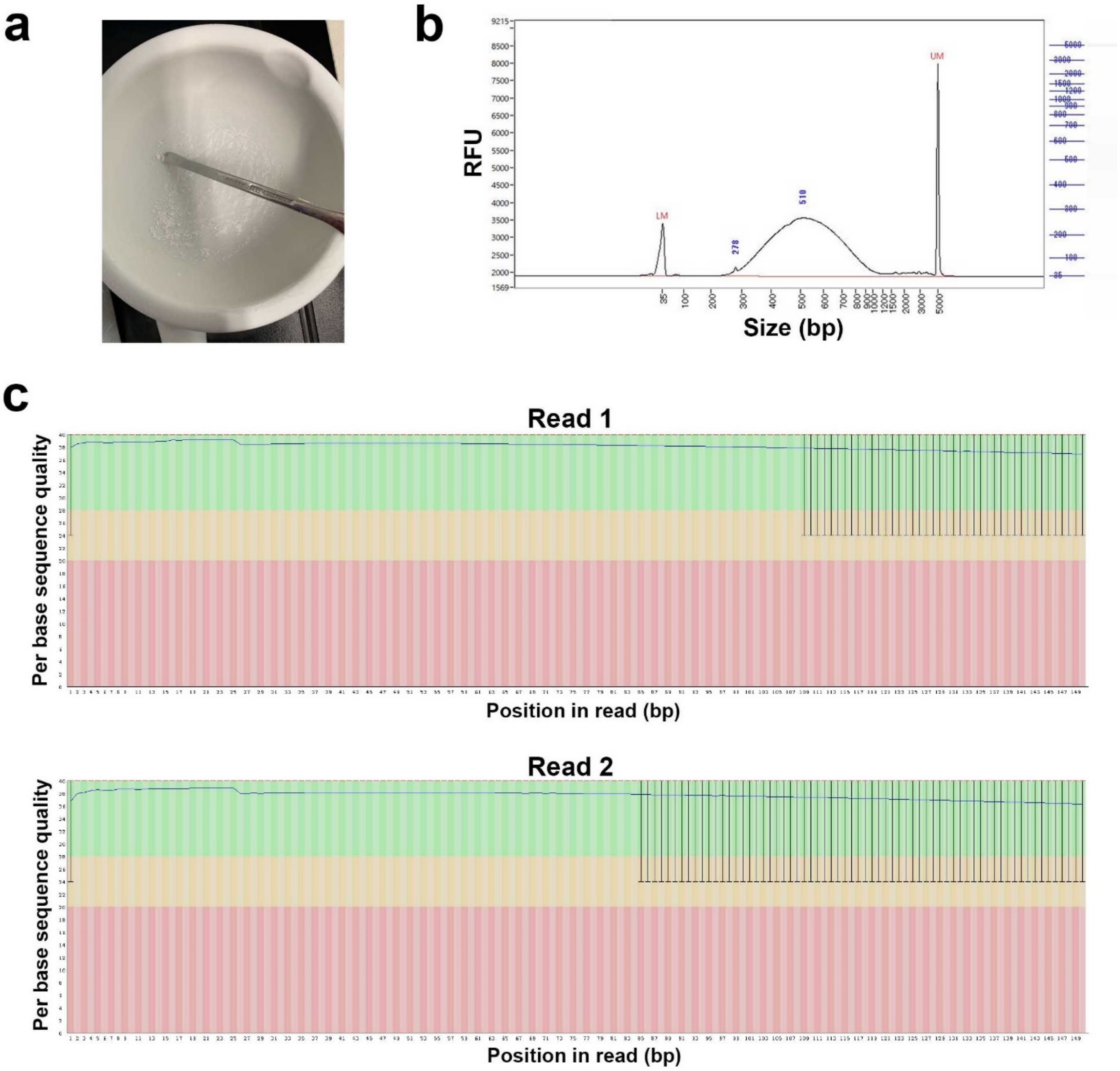


**Fig. S5. Hi-C sequencing of the *G. rufa* genome**

(a) Flash-frozen tissue ground into a fine powder. (b) Insert size distribution of the Hi-C library measured by Bioanalyzer (Agilent, CA, USA). The result is shown in the electropherogram (left panel) and gel image (right panel). The insert size distribution is approximately between 300 bp and 1,000 bp, meeting the quality requirements specified in the manufacturer's protocol of the Dovetail Omni-C Kit (Cantata Bio, CA, USA). Abbreviations: LM, lower marker; RFU, relative fluorescence unit. (c) Sequence quality of the Hi-C sequencing reads.

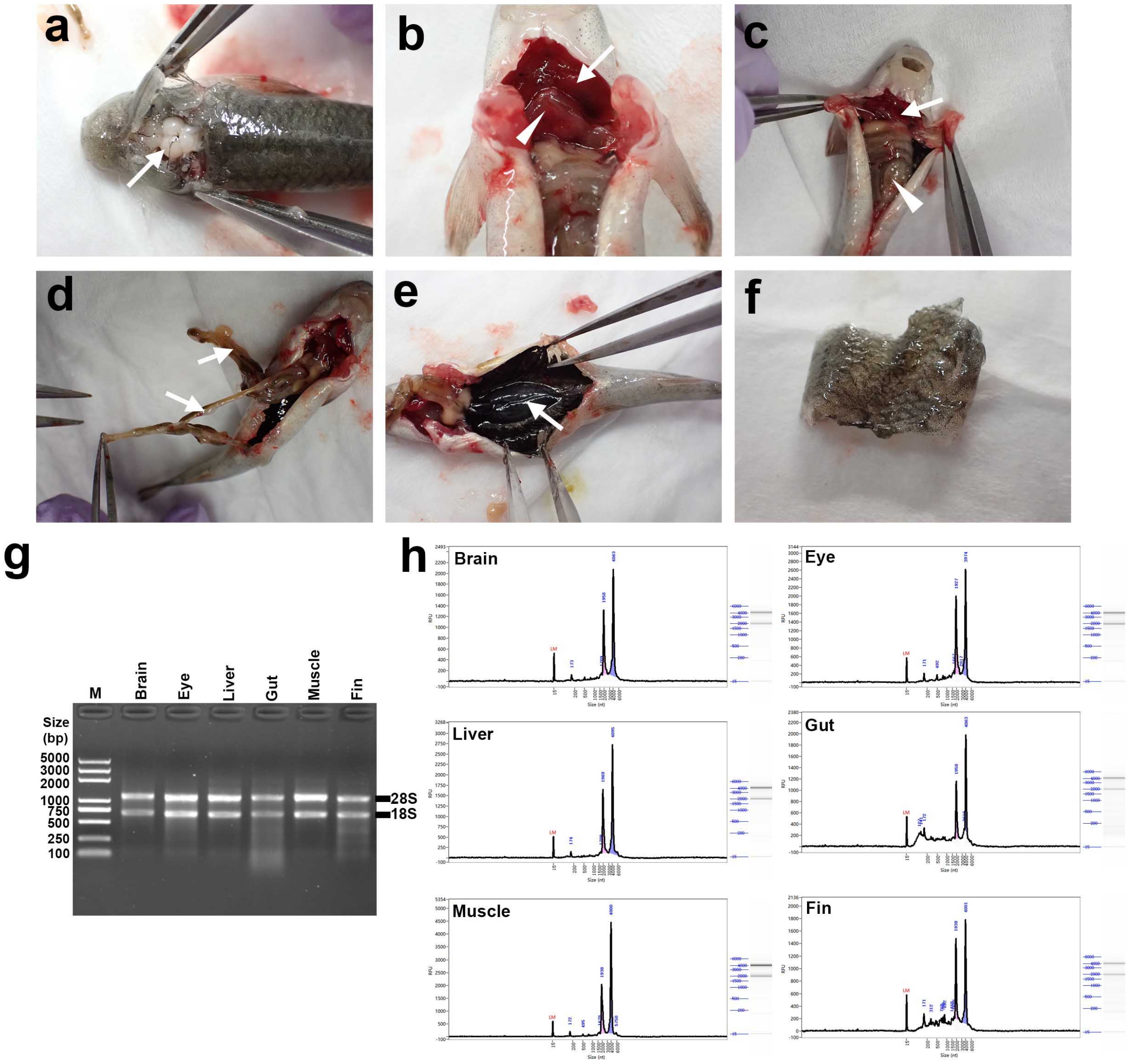


**Fig. S6. Total RNA extraction from *G. rufa* tissues for RNA-seq analysis**

(a) Macroscopic view after opening the skull, showing the brain (white arrow). (b) Macroscopic view after opening the abdominal wall, showing the gills (white arrow) and the heart (arrowhead). (c) Macroscopic view showing the liver (white arrow) and gut (white arrowhead). (d) Macroscopic view after cutting the peritoneum and pulling the gut (white arrows) outside the body. (e) Retroperitoneal region, indicating the kidney area (white arrows). (f) Dissected skin tissue. (g) Agarose gel electrophoresis of extracted total RNA. Agarose gel electrophoresis of extracted total RNA was performed using a 1% agarose gel, with a voltage of 180 V and a run time of 16 min. M represents the molecular weight marker. Note that the 28S and 18S ribosomal RNA bands are clearly visible, suggesting that the total RNA is minimally degraded. (h) Size distribution of the extracted total RNA measured using the Agilent 5400 Fragment Analyzer System (Agilent, CA, USA). The horizontal axis represents size and the vertical axis represents relative fluorescence unit (RFU). Abbreviations: LM, lower marker.

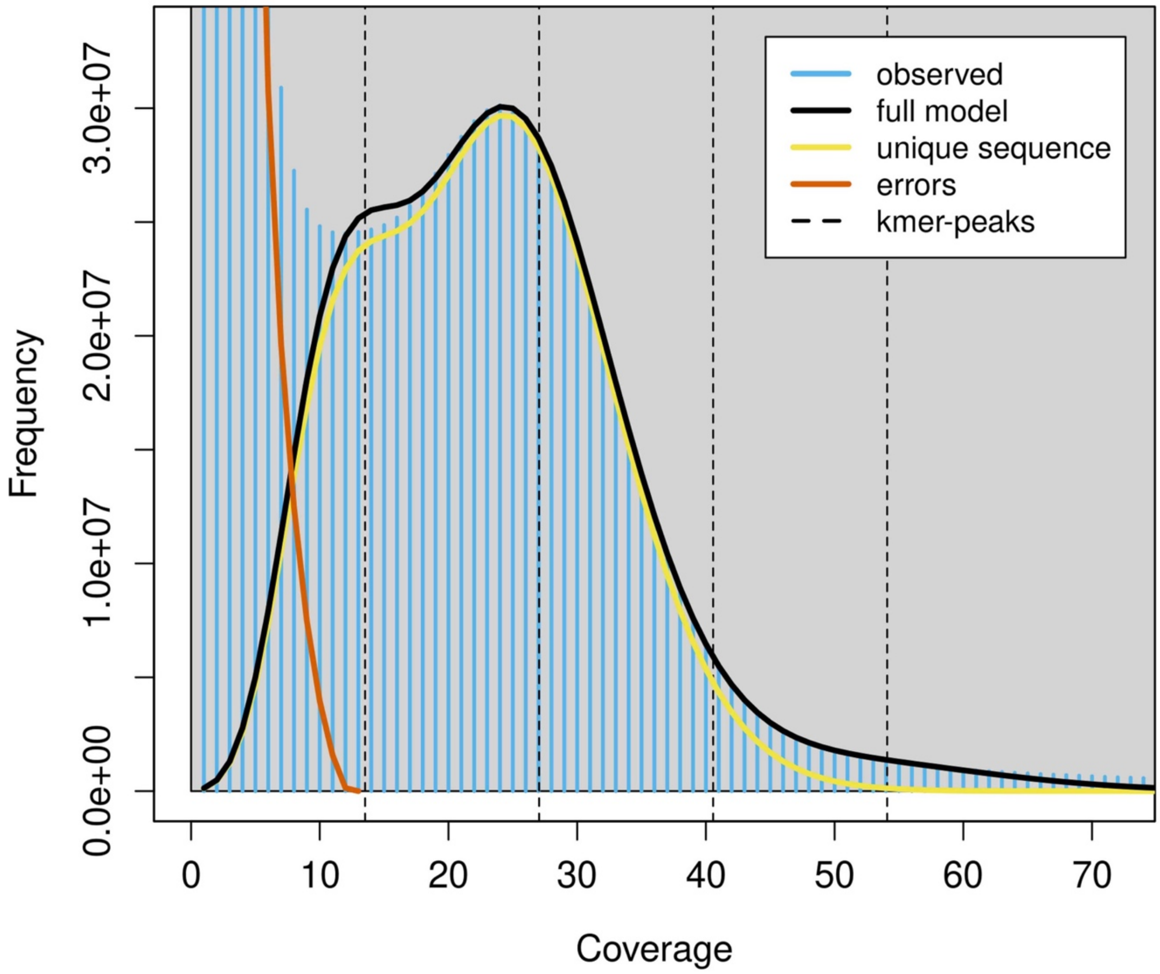


**Fig. S7. The k-mer distribution of the Illumina genome sequencing reads using GenomeScope based on a k value of 21**

The horizontal axis represents the k-mer coverage and the vertical axis represents the frequency of the k-mer for a given coverage. The first and second peaks are composed of the homozygous and heterozygous contents, respectively. The estimated genome size was 1.14 Gb, and the estimated heterozygosity was 1.03%. The estimated genome coverage of the repetitive elements was 45.4%.

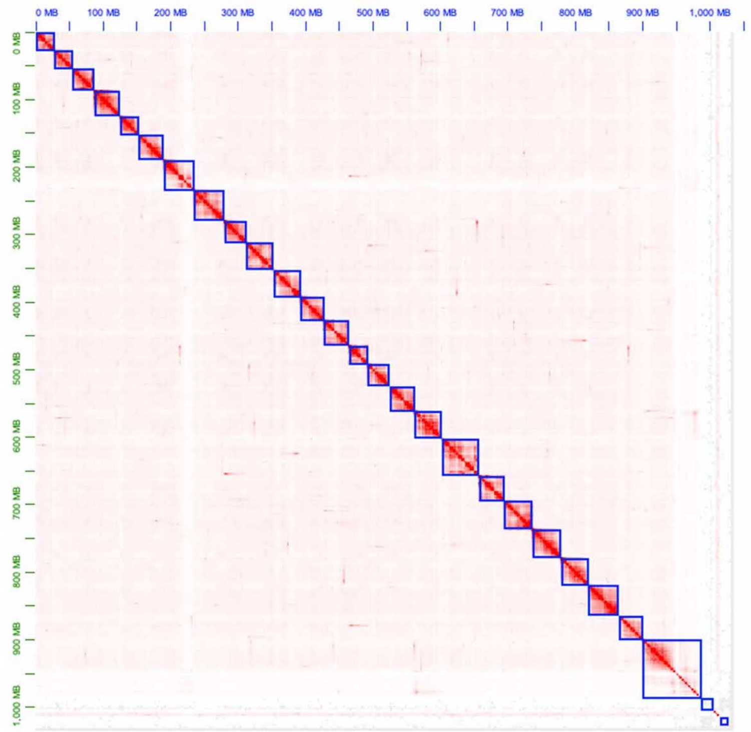


**Fig. S8. Hi-C contact map of the haplotype 2**

An Hi-C contact map of haplotype 2 is shown. Each blue square represents the boundary between the scaffolds. Chromatin contact signals are shown as red spots, and their intensities reflect the degree of chromatin contact strength.

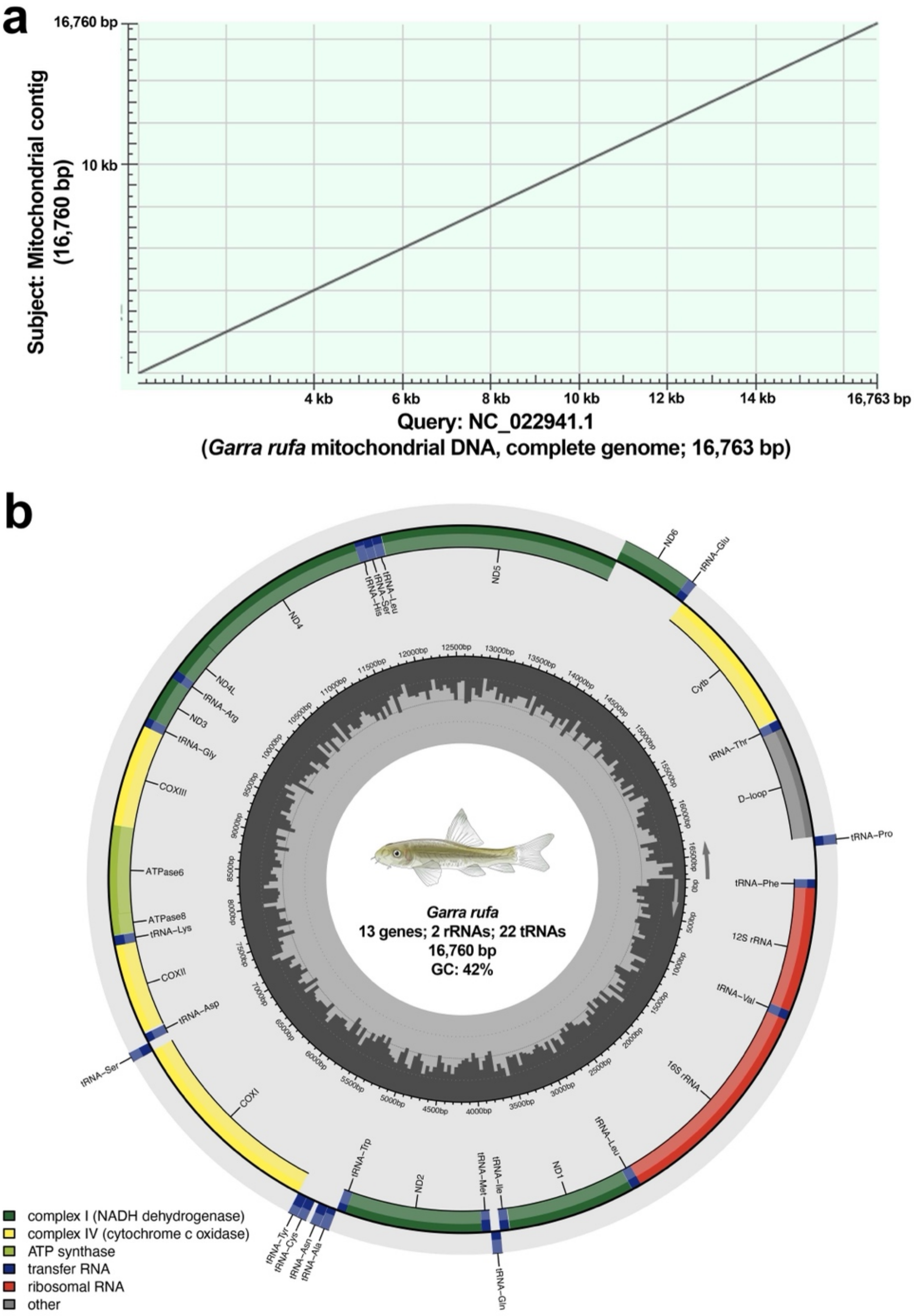


**Fig. S9. Mitochondrial genome of *G. rufa***

(a) Alignment between the publicly available mitochondrial genome sequence of *G. rufa* (NC_022941.1) and the mitochondrial contig assembled in the present study. An alignment was obtained across the full length of both sequences, showing 99.24% sequence identity with two gaps. This figure was generated using the NCBI BLAST web server with the default parameters. (b) The circular mitochondrial genome is described with genes shown by colored blocks. Genes located inside the map are on the forward strand, while those outside are on the reverse strand. The D-loop is shown in grey, the ribosomal RNA genes (16S and 12S) are shown in red, the 22 transfer RNA genes are shown in blue, and the 13 protein-coding genes are shown in dark green, light green, and yellow. The innermost circle represents the GC content of the regions.

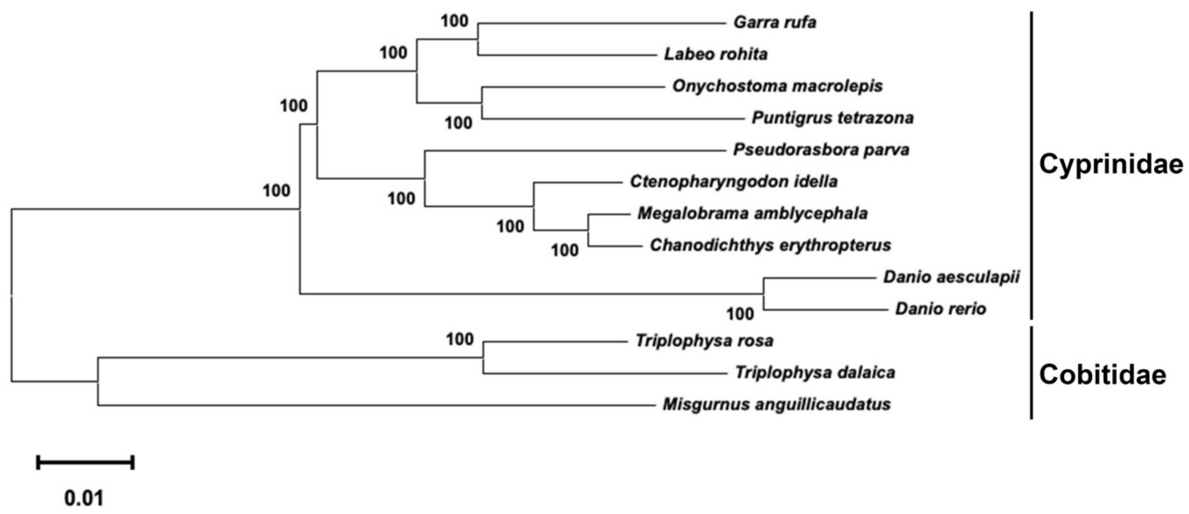


**Fig. S10. Neighbor-joining tree showing the relationship between *G. rufa* and other cypriniforms**

A neighbor-joining tree was constructed to show the phylogenetic relationships between *G. rufa* and the 12 Cypriniformes species. The scale bar represents the phylogenetic distance of 0.01 nucleotide substitutions per site.
